# Public volume electron microscopy data: An essential resource to study the brain microvasculature

**DOI:** 10.1101/2022.02.20.481154

**Authors:** Stephanie K. Bonney, Vanessa Coelho-Santos, Sheng-Fu Huang, Marc Takeno, Joergen Kornfeld, Annika Keller, Andy Y. Shih

## Abstract

Electron microscopy is the primary approach to study ultrastructural features of the cerebrovasculature. However, 2D snapshots of a vascular bed capture only a small fraction of its complexity. Recent efforts to synaptically map neuronal circuitry using volume electron microscopy have also sampled the brain microvasculature in 3D. Here, we perform a meta-analysis of 6 data sets spanning different species and brain regions, including 2 data sets from the MICrONS consortium that have made efforts to segment vasculature in addition to all parenchymal cell types in mouse visual cortex. Exploration of these data have revealed rich information for detailed investigation of the cerebrovasculature. Neurovascular unit cell types (including, but not limited to, endothelial cells, mural cells, perivascular fibroblasts, microglia, and astrocytes) could be discerned across broad microvascular zones. Image contrast was sufficient to identify subcellular details, including endothelial junctions, caveolae, peg-and-socket interactions, mitochondria, Golgi cisternae, microvilli and other cellular protrusions of potential significance to vascular signaling. Additionally, noncellular structures including the basement membrane and perivascular spaces were visible and could be traced between arterio-venous zones along the vascular wall. These explorations revealed structural features that may be important for vascular functions, such as blood-brain barrier integrity, blood flow control, brain clearance, and bioenergetics. They also identified limitations where accuracy and consistency of segmentation could be further honed by future efforts. The purpose of this article is to introduce these valuable community resources within the framework of cerebrovascular research by providing an assessment of their vascular contents, identifying features of significance for further study, and discussing next step ideas for refining vascular segmentation and analysis.

## INTRODUCTION

Electron microscopy is an essential tool in cerebrovascular research. It is the primary means to visualize and quantify subcellular structures central to blood-brain barrier function, blood flow regulation and neurovascular communication. These functions rely on endothelial tight and adherens junctions, caveolar vesicles, organelles (e.g. mitochondria), and the vascular basement membrane, for example, which are all structures on the order of tens to hundreds of nanometers. Two-dimensional (2D) transmission electron microscopy (TEM) is most often used to resolve these ultrastructures, providing cross sectional high-resolution views of the vascular wall. However, vascular cells and their subcellular compartments have sophisticated 3D morphologies inadequately captured in 2D images. For example, mural cells (smooth muscle cells and pericytes) exhibit diverse cellular morphologies and cellular interactions with the endothelium in different regions of the microvasculature (Grant et al., 2017;Ornelas et al., 2021). “Peg-and-socket” contacts between pericytes and endothelial cells, sites where the direct communication between these cells takes place, can only be visualized using electron microscopy. These contacts are likely important for cerebrovascular function since loss of pericyte-endothelial interactions are associated with increased blood-brain barrier permeability (Armulik et al., 2010;Daneman et al., 2010), brain entry of circulating leukocytes (Török et al.), and impaired cerebral blood flow (Kisler et al., 2017;Nikolakopoulou et al., 2019;Hartmann et al., 2021a).

A second limitation of non-automated 2D EM is that only fields of view in the micrometer range can be imaged, a scale on which microvasculature is sparse. Further, there are distinct functional zones of the microvasculature, including arterioles, capillaries, venules or transitional regions between them, that are difficult to locate and image in 2D. The cellular composition of the vascular wall, *i.e.* the neurovascular unit, differs between the vascular zones. Arterioles are composed of endothelial cells, smooth muscle cells, perivascular fibroblasts and macrophages, and astrocytic endfeet. In contrast, capillaries are composed of endothelial cells, pericytes encased in the basement membrane and astrocytic endfeet. The subcellular structures and organelles of these cell types and the composition of the basement membrane between the cells also differ between microvascular regions. Therefore, to understand the vasculature, data must be collected at an ultrastructural level on a scale of hundreds of micrometers to millimeters.

Recent advances in 3D-EM of brain tissue have overcome these considerable technical challenges (Kornfeld and Denk, 2018;Yin et al., 2020), and several high-content 3D-EM data sets from various species have been generated (e.g. finch, mouse, human). The primary drive behind these efforts has been to map synaptic connectivity between neurons. Initial data sets, such as the *C. elegans* data captured in the 1980s (White et al., 1986), were of smaller size due to challenges in imaging, segmentation and data processing. However, recent collaborative efforts through the Machine Intelligence from Cortical networks (MICrONS) consortium (Dorkenwald et al., 2019;Consortium et al., 2021;Schneider-Mizell et al., 2021) and independent laboratories (Lee et al., 2016;Dorkenwald et al., 2017;Bloss et al., 2018;Shapson-Coe et al., 2021) have generated enormous data sets encompassing up to a cubic millimeter of tissue volume. These publicly available 3D-EM data sets hold immense information on the fine-structure of cerebral microvasculature across different zones of the vascular network, allowing deep exploration of cellular composition, morphology and subcellular interaction between neurovascular cell types. Despite being available in online browsers, these data have yet to be mined and interpreted for insight on cerebrovascular biology. The purpose of this article is to assess large-scale 3D-EM data sets gathered from the brain, and to show vignettes of data on the cerebral vasculature held within them.

## METHODS

All data sets were explored through the access links provided in Table 1. These resources have used Neuroglancer, an open-source browser-based viewer for visualization of large-scale 3D data. An introduction on how to use Neuroglancer can be found at: https://www.microns-explorer.org/visualization.

**Table 1.**
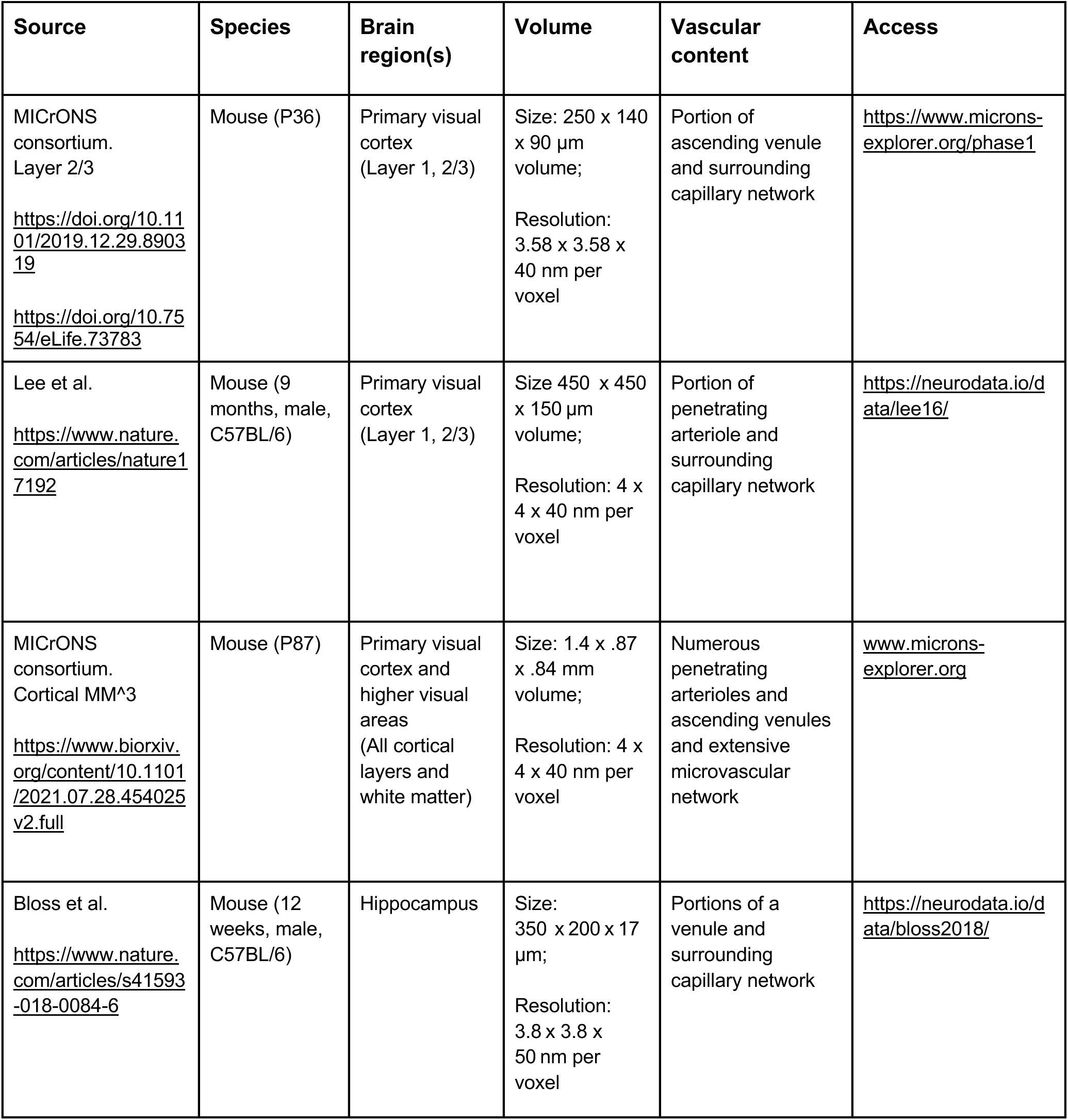
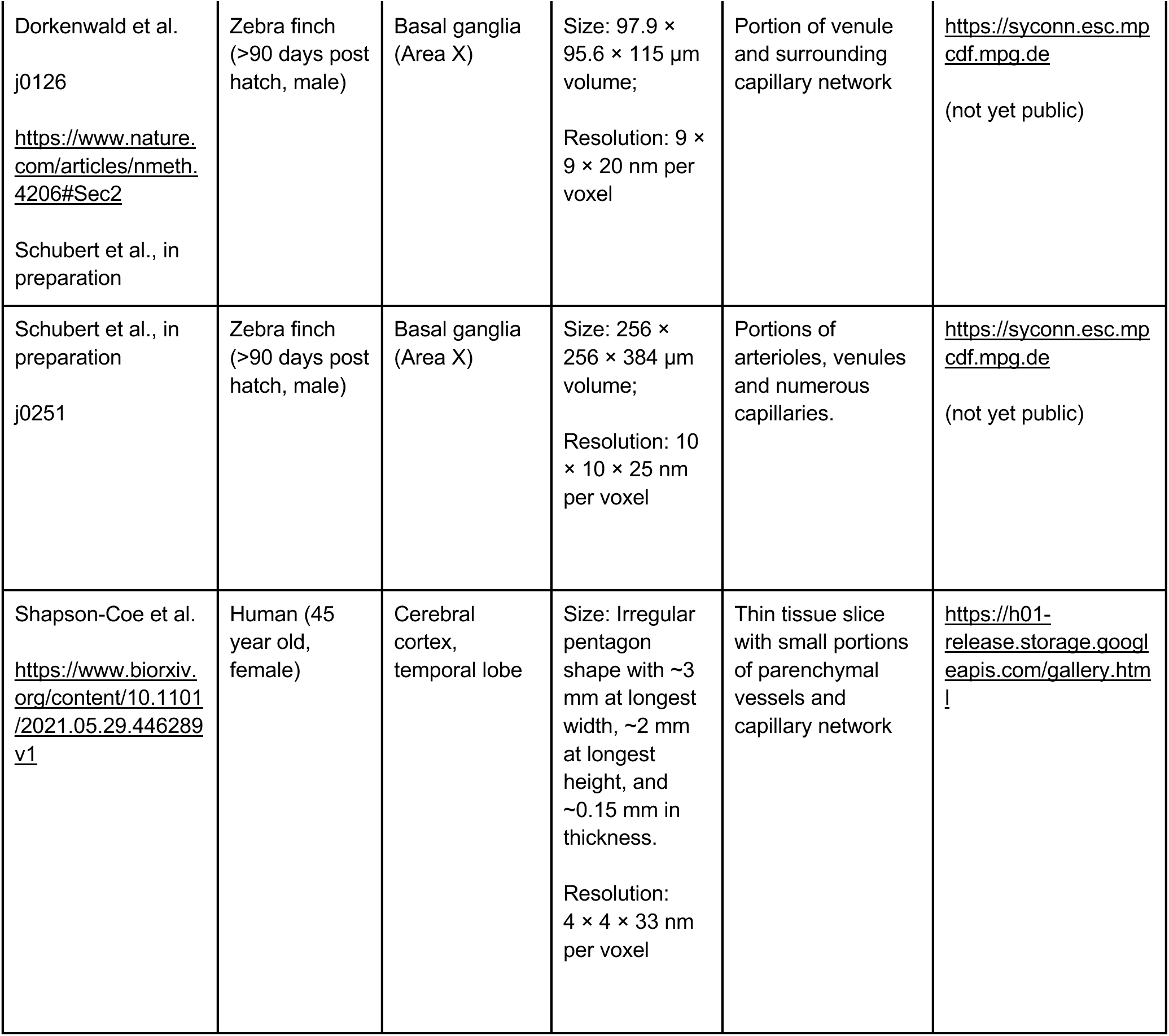
Attributes and vascular contents in public 3D-EM data sets from brain tissue.

Full details on Neuroglancer can be found at: https://github.com/google/neuroglancer#readme.

In the figure legends, we provide the x,y,z coordinates for the regions of interest shown in each data set, which can be copied and pasted into the x,y,z, query boxes on the top left of Neuroglancer.

For Figures 1 and 3, web addresses are provided in the figure legend to view annotated MICrONS Layer 2/3 data. However, viewers must first register and agree to terms of service, which can be prompted through the following link: https://neuromancer-seung-import.appspot.com/?json_url=https://globalv1.daf-apis.com/nglstate/api/v1/5665719098277888.

The data sets examined here are not exhaustive of those available for public viewing. The reader is directed to the following websites for additional volume EM resources: https://neurodata.io/project/ocp/. https://webknossos.org/publications.

## RESULTS

**Table 1** provides an overview of the publicly available 3D-EM data sets examined in this study. We first focus on two data sets from the cerebral cortex of adult mice, generated by the MICrONS consortium, which can be accessed through www.microns-explorer.com.

### MICrONS Layer 2/3

An initial study examined a region of adult mouse visual cortex spanning a volume of 250 x 140 x 90 µm in cortical layers 2/3 (Schneider-Mizell et al., 2021). All parenchymal cell types were segmented in this data set (neurons, astrocytes, microglia) and the vascular wall was segmented as a combined element of both mural cells and endothelial cells. The vascular network includes a portion of a single cortical ascending venule and surrounding capillaries. Occasionally, neurons or astrocytes are captured within the vascular segmentation due to their proximity to the vascular wall.

#### Pericyte-endothelial interaction

We noted that the capillary network contains 25 endothelial cell nuclei and 5 pericyte nuclei, consistent with individual pericytes covering multiple endothelial cells (**Fig. 1A**). Pericytes were identified as cells on the abluminal side of endothelium with protruding cell somata and elongated, slender processes embedded in the vascular basement membrane. In the volume EM data, their abluminal processes could be traced back to the cell somata to verify its identity as a pericyte. Physical interlocking between pericytes and endothelial cells via peg-and-socket interactions is important for their communication and attachment. These structures could be discerned in the data (**Fig. 1B**). By annotating the positions of peg-and-socket interactions in Neuroglancer, they were found to be concentrated near the somata of pericytes, implicating the pericyte somata as potential hot spots for direct communication with endothelial cells (**Fig. 1B**, C). These interactions included both extension of pericytes pegs toward the adjacent endothelium, and less commonly, endothelial cells extending toward pericytes (**Fig. 1B, D**).

**Figure 1.**
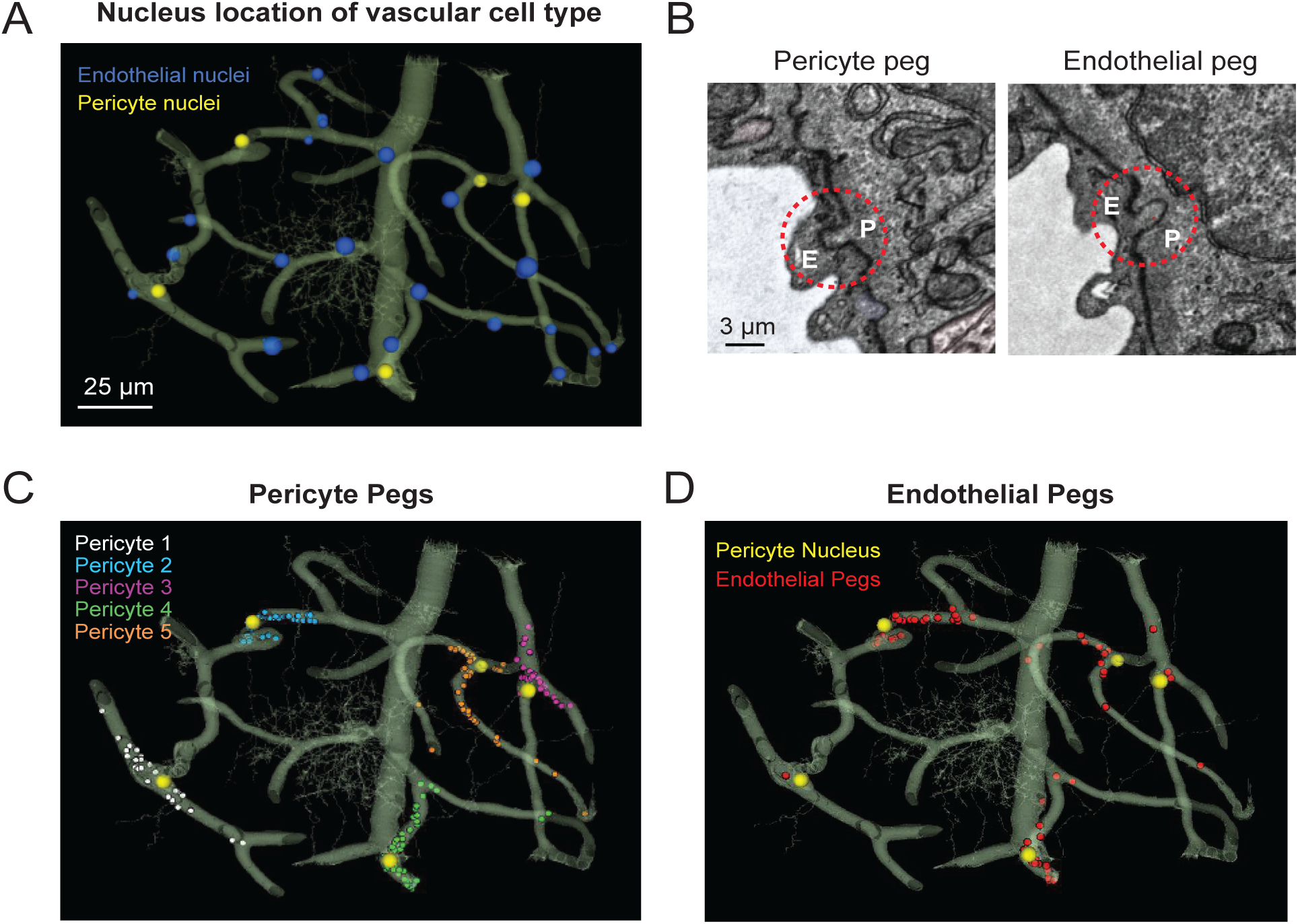
MICrONS Layer 2/3 pericyte-endothelial interactions. **A.** The location of all endothelial and pericyte nuclei identified within the MICrONS Layer 2/3 data set. **B.** Examples of pericyte pegs extending into the endothelium (left), and conversely, endothelial pegs extending into the pericyte (right). **C.** The location of pericyte pegs mapped onto the 3D microvascular network reveals that pegs are enriched at sites of pericyte somata. The pericyte somata are labeled with yellow spheres, and the individual pegs are labeled with smaller colored spheres, color-coded per pericyte. **D.** Endothelial pegs are also enriched around pericyte somata. Images adapted from (Ornelas et al., 2021). Link to annotations (See methods to register and agree to terms of service): https://neuromancer-seung-import.appspot.com/?json_url=https://globalv1.daf-apis.com/nglstate/api/v1/5073246747623424.

Pericyte somata are often positioned at capillary bifurcations (**Fig. 2A**)(Hartmann et al., 2015). This localization may play a role in how blood flow is partitioned down separate capillary branches during cerebral perfusion (Gonzales et al., 2020). Endothelial cell nuclei were located along capillary segments and also some capillary bifurcations (**Fig. 2B**). The extra volume of the endothelial nuclei often manifested as abluminally-oriented protrusions, similar to pericyte somata (**Fig. 2C**). However, we also noted many instances where the endothelial nuclei encroached upon the intraluminal space, reducing its diameter (**Fig. 2D**). This suggests that the position of endothelial nuclei may influence blood cell flow in the capillary network by imparting blood flow resistance. This possibility has not been thoroughly examined in physiological studies, unlike blood flow in relation to the position of pericyte somata.

**Figure 2.**
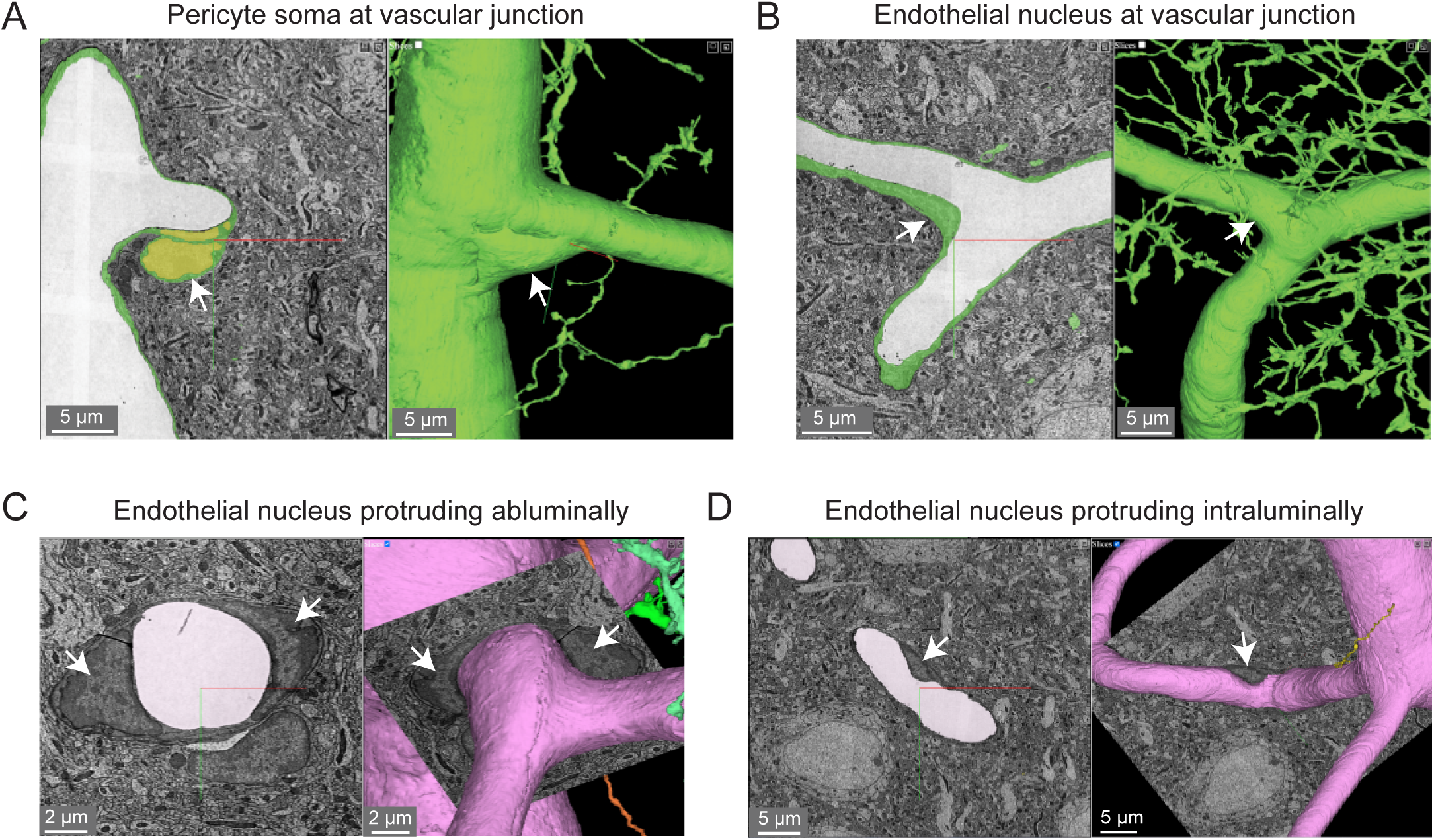
Pericyte and endothelial cell somata can be positioned at vascular junctions. **A.** Pericyte soma (white arrow) positioned at the junction between the capillary and ascending venule in the MICrONS Layer 2/3 data set. Left panel shows selected vascular wall segmentation in 2D, and the right panel shows 3D rendering of the vascular wall. MICrONS Layer 2/3 x,y,z coordinates at: 82286, 68574, 1719. **B.** Endothelial nucleus (white arrow) at a capillary junction. Coordinates at: 68291, 61335, 1788. **C.** Example of two endothelial nuclei that protrude abluminally without affecting the shape of the lumen. Coordinates at: 76691, 59357, 1594. **D.** Example of an endothelial nucleus that protrudes intraluminally and causes local reduction in capillary diameter. Coordinates at: 71372, 45504, 1467.

The majority of the capillary length in cortex (∼90%) is contacted by the slender processes of pericytes (Grant et al., 2017;Berthiaume et al., 2018). To better understand the arrangement of pericyte processes in specific regions of interest, we manually labeled their edges with sequential annotations in Neuroglancer (red dots; **Fig. 3A, Ai**). This allowed us to track the pericyte process alongside co-annotated endothelial features (somata, cell-cell junctions)(Ornelas et al., 2021). Pericyte processes were occasionally found to cover and track along with endothelial cell junctions, suggesting that they may provide structural support for this feature of the blood-brain barrier. When pericyte processes encountered the location of endothelial nuclei, they tended to increase their surface coverage and proximity to the nucleus (**Fig. 3B, Bi**)(Ornelas et al., 2021). Heightened coverage of the endothelial nucleus may provide enhanced communication that influence nuclear functions, such as gene expression.

**Figure 3.**
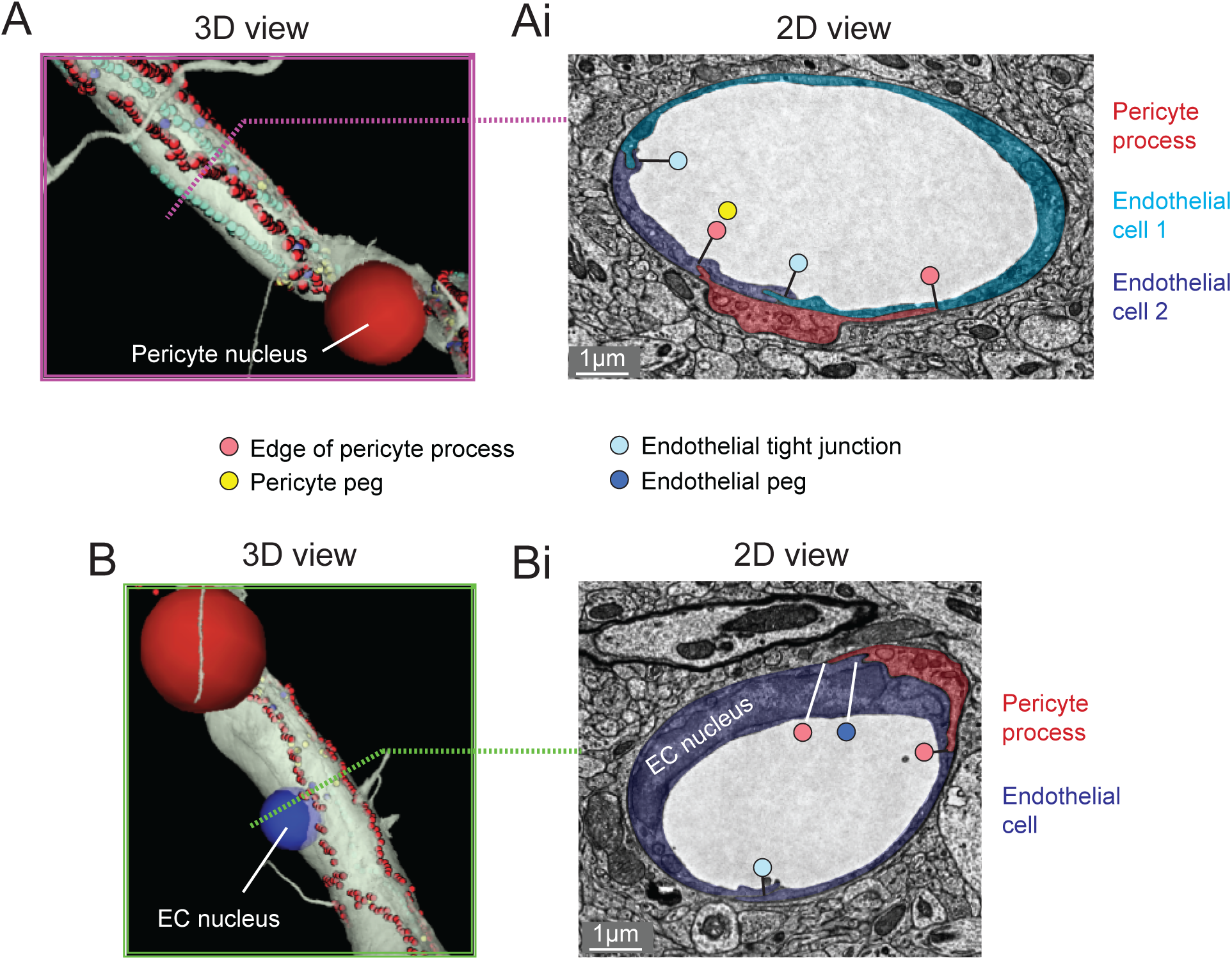
Pericyte processes in relation to endothelial features. **A.** The edge of a pericyte process annotated in Neuroglancer 3D view with red dots, alongside endothelial junctions in cyan dots. **Ai.** The 2D cross section of one location (dotted magenta line) within the same capillary segment. **B.** The pericyte process tends to exhibit greater endothelial coverage near endothelial somata. **Bi.** The 2D cross section of one location (dotted green line) within the same capillary segment. EC = endothelial cell. Images adapted from (Ornelas et al., 2021). Link to annotations (See methods to register and agree to terms of service): https://neuromancer-seung-import.appspot.com/?json_url=https://globalv1.daf-apis.com/nglstate/api/v1/6082177280245760.

#### Endothelial protrusions

“Microvilli” are small protrusions that extend from the endothelium into the intraluminal space (**Fig. 4A**). They have been widely observed across mammals and birds (Fujimoto et al., 1975;Makarov et al., 2015), though their physiological role has remained poorly understood. They could be important for sensing the passage of blood cells or impede the movement of blood cells by creating resistance. Indeed, prior studies have shown that microvilli incidence increases in cerebral ischemia, perhaps contributing to poor capillary blood flow and no reflow (Dietrich et al., 1984). 3D-EM data makes it possible to study the shape, prevalence and distribution of microvilli throughout the capillary network. Curiously, there were also some instances where the endothelium protruded abluminally toward the brain parenchyma (**Fig. 4B**). Perhaps these are also specialized appendages for direct sensation of neural or astrocytic activity. In fact, the protrusion shown is surrounded by neurites and one axon swells into a bouton shape when in contact with the endothelium (**Fig. 4C**).

**Figure 4.**
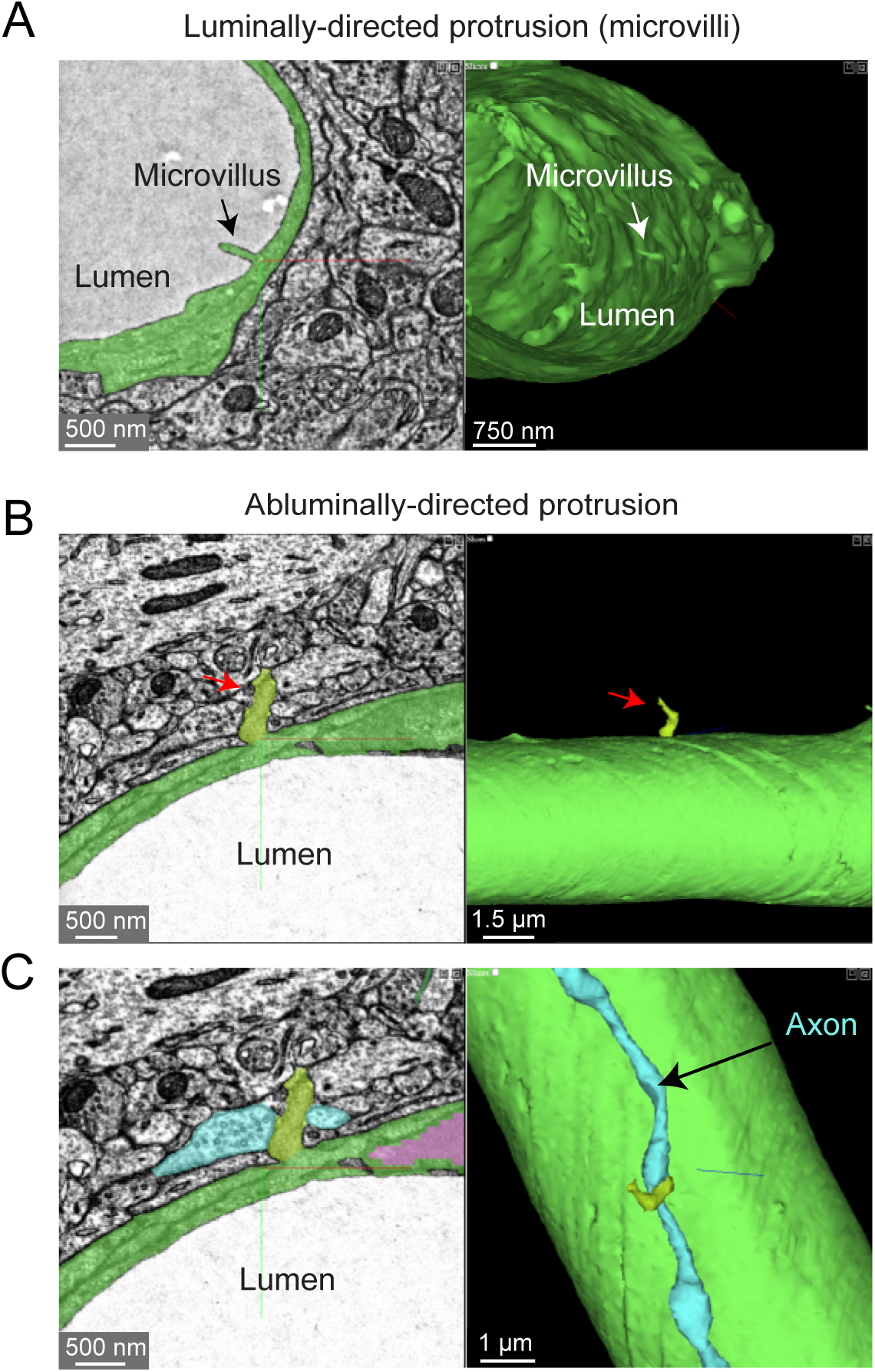
Endothelial protrusions extend luminally and abluminally from the vessel wall. **A.** Example of a luminally-directed endothelial protrusion (microvilli) within a capillary segment. Left panel shows the 2D cross section of the structure (black arrow). Right panel shows a 3D view of the same structure in segmented vasculature (white arrow). MICrONS Layer 2/3 x,y,z coordinates at: 62431, 48503, 1336. **B.** Example of abluminally-directed endothelial protrusion. Left panel shows the 2D cross-section of the structure (black arrow). Right panel shows a 3D view of the same structure in segmented vasculature. MICrONS Layer 2/3 x,y,z coordinates at: 73675, 48870, 746. The structure is segmented separately from the vascular wall, but is in fact an extension of the endothelial cell. **C.** The abluminally-directed endothelial protrusion is surrounded by neuronal axons and dendrites. One axon (cyan) exhibits a region of swelling (potential bouton) in direct contact with the endothelial structure.

#### Astrocyte coverage

Astrocytes are extremely complex in structure and their multitude of fine processes interact with the vasculature in the form of “endfeet”, which cover much of the abluminal surface (Mathiisen et al., 2010;Korogod et al., 2015). Astrocytic coverage of the vasculature contributes to blood-brain barrier integrity (Araya et al., 2008), release of vasoactive substances (Attwell et al., 2010), and water transport (Jin et al., 2013). The segmented astrocytes in the MICrONS data sets have classic astrocyte morphology (dense and highly ramified process) with endfeet in subregions of the cell that covered the vascular wall. We observed that astrocytes covered the vasculature in discrete non-overlapping territories (**Fig. 5A**). Along just one capillary segment, multiple astrocytes cooperated to establish coverage (Kubotera et al.). However, it is difficult to know if individual astrocytes have been wholly segmented, given their morphological complexity. Thus, it is also possible that the image shows fragments of individual astrocytes. These data may also be useful to understand the extent and distribution of regions that lack astrocyte coverage, which may be relevant as sites of immune cell entry and interaction (Horng et al., 2017).

**Figure 5.**
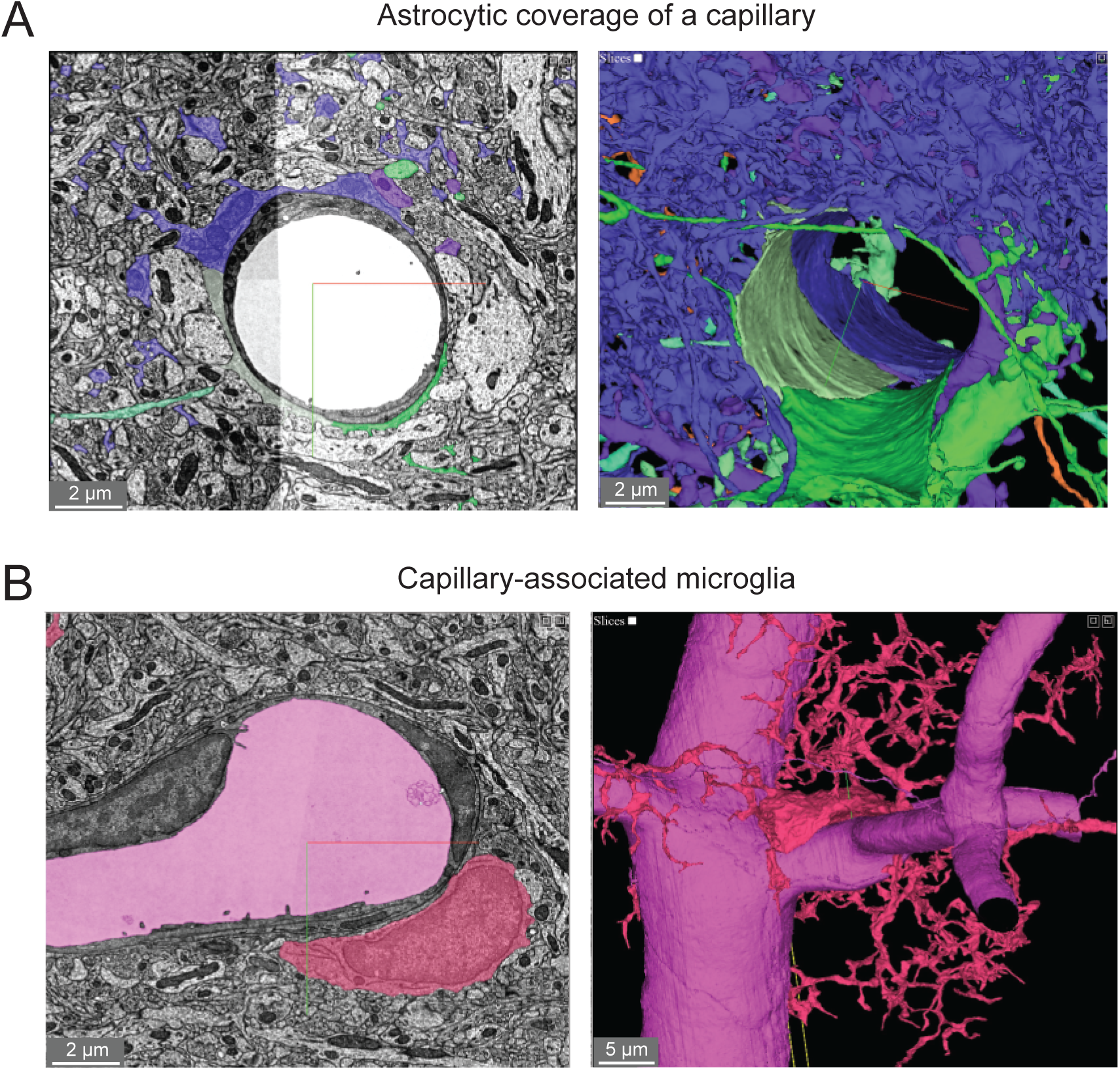
Astroglial and microglial vascular interactions. **A.** Three astrocytes (blue, green, light green) enwrap the capillary wall and their territories form a tiling pattern. Dendrites of a neuron (purple) course through the astrocytic territory. The vascular wall is not shown in the 3D view (right) to provide a clearer view of astrocyte contact with the vascular wall. MICrONS Layer 2/3 x,y,z coordinates at: 77091, 62413, 784. **B.** A putative capillary-associated microglia. The 3D view shows the vascular lumen in fuchsia and a microglial soma (red) that drapes along the vessel surface at a bifurcation. Close inspection of this microglia will reveal that it directly contacts the vascular basement membrane without intervening astrocytic endfeet. MICrONS Layer 2/3 x,y,z coordinates at: 76302, 59416, 1641.

#### Microglial-capillary interaction

Microglia associate with the capillary wall and contribute to regulation of neurovascular structure and function (Joost et al., 2019;Bisht et al., 2021). Capillary-associated microglia could be detected in the data set and close inspection revealed that they abutted the vascular basement membrane without intervening contact with astrocytic endfeet (**Fig. 5B**), consistent with prior studies (Mathiisen et al., 2010;Bisht et al., 2021). Individual microglia are segmented allowing for detailed structure and ultrastructural investigation of their interactions with other neurovascular cells.

#### Subcellular features

Two subcellular features are often quantified in studies that assess blood-brain barrier integrity. First are electron dense tight junctions where individual endothelial cells are bound by a variety of adhesion proteins to form a barrier (claudins, occludin, junction adhesion molecules, cadherins)(Haseloff et al., 2015). Endothelial cell-cell junctions are not simply rigid physical barriers, but highly dynamic structures regulating the paracellular (between cell) passage of cells, water, ions and macromolecules. Second, small endothelial vesicles called caveolae facilitate transcellular (through cell) passage across the endothelium (Andreone et al., 2017). Both of these classic blood-brain barrier structures are clearly visualized in the Cortical Layer 2/3 data set (**Fig. 6**).

**Figure 6.**
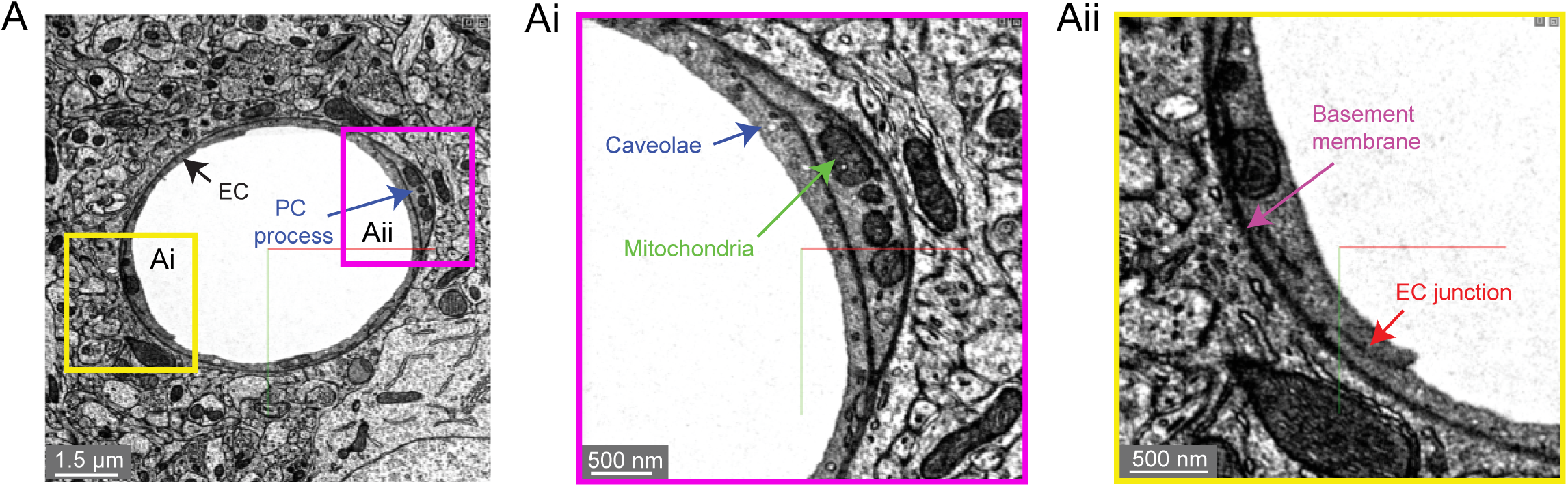
Subcellular structures in the vasculature are visible with high contrast in the MICrONS Layer 2/3 data set. **A.** A representative cross-section of a capillary with magnified insets showing commonly measured subcellular structures, including caveolae and mitochondria (**Ai**) and endothelial junctions and basement membrane (**Aii**). MICrONS Layer 2/3 x,y,z coordinates at: 109802, 58170, 707.

We noticed that caveolae density varied among capillary segments, which may mean that transcellular permeability occurs heterogeneously across the capillary network (not shown). 3D-EM data may potentially allow caveolae density and their characteristics to be charted across the microvascular network to better understand their distribution. Recent studies have already highlighted interesting zone-specific differences that involve caveolae. Venous endothelial cells and capillaries exhibit greater uptake of plasma proteins through transcytosis than arterioles (Yang et al., 2020). Similarly, post-capillary venules show heightened receptor-mediated transport of nanoparticle useful for drug delivery (Kucharz et al.). Interestingly, arteriole endothelial cells are not devoid of caveolae and in fact show an enrichment in caveolae essential to neurovascular signaling during functional hyperemia (Chow et al., 2020). Mitochondria are segmented throughout the data set, including in endothelial cells and mural cells (**Fig. 6A, Ai**)(Turner et al., 2020). Prior studies have shown pericytes to be enriched with mitochondria (Mathiisen et al., 2010), which are likely needed to support the metabolic demands of solute transport at the blood-brain barrier.

#### Non-cellular features

An important component of the vascular wall is the basement membrane (BM) between cells that contribute to cellular adhesion and signaling. The basement membranes are thin and highly organized structures (50-100 nm in thickness) consisting mainly of collagen IV, fibronectin, laminins, nidogen and proteoglycan perlecan (Xu et al.). There can be multiple BM layers depending on the microvascular zone. In capillaries, there is the endothelial BM that lies between the endothelium and the pericyte, and the parenchymal BM between the pericyte and the astrocyte. The basement membrane is very apparent in 3D-EM data because it is generally more electron dense than the neighboring cells (**Fig. 6A, Aii**). At sites of pericyte-endothelial interaction, the basement membrane may become thin and harder to recognize. Increased basement membrane thickness is described in aging and conditions of poor vascular health such as in the tissues surrounding an ischemic stroke (Nahirney et al., 2015). This may indicate a loosening of the protein sheets or adverse production of BM proteins, both of which would reduce neurovascular communication.

### MICrONS Cortical MM^3

A second study from the MICrONS consortium imaged a 1.4 x .87 x .84 mm volume from visual cortex, which is 400x larger than the prior Layer 2/3 sample. It spans all 6 cortical layers, including a small amount of white matter at its base. This significantly larger tissue volume provides the unique opportunity to study the neurovascular unit across different vascular zones, cortical layers, multiple cortical areas and between gray versus white matter.

#### Microvascular architecture

We first briefly discuss the vascular architecture of the cerebral cortex. Blood flow to the cortex enters through leptomeningeal arterioles on the brain surface. From the pial arteriolar network, penetrating arterioles branch and dive into the parenchyma to perfuse columns of tissue (Shih et al., 2015). Penetrating arterioles then send smaller offshoots called arteriole-capillary transitions (or pre-capillary arterioles) that successively ramify into dense and highly interconnected capillary networks. After passing the dense capillary network, blood then coalesces into ascending venules oriented in parallel with penetrating arterioles, that drain back to a network of leptomeningeal venules at the pial surface.

The vascular network of the MICrONS Cortical MM^3 data set can be visualized in entirety by selecting the segmentation of the intraluminal space. The data contains leptomeningeal vessels within the subarachnoid space of the meninges, which can be followed as they dive into penetrating vessels of the cortical parenchyma. In total, 12 penetrating arterioles and 25 ascending venules were captured within the volume (**Fig. 7A**). The identity of the penetrating vessel was determined by examining the morphology of mural cells that lie abluminal to the endothelium. Arterioles are surrounded by concentric, ring-shaped smooth muscle cells (**Fig. 7B**), whereas venules lack these cells (**Fig. 7C**).

**Figure 7.**
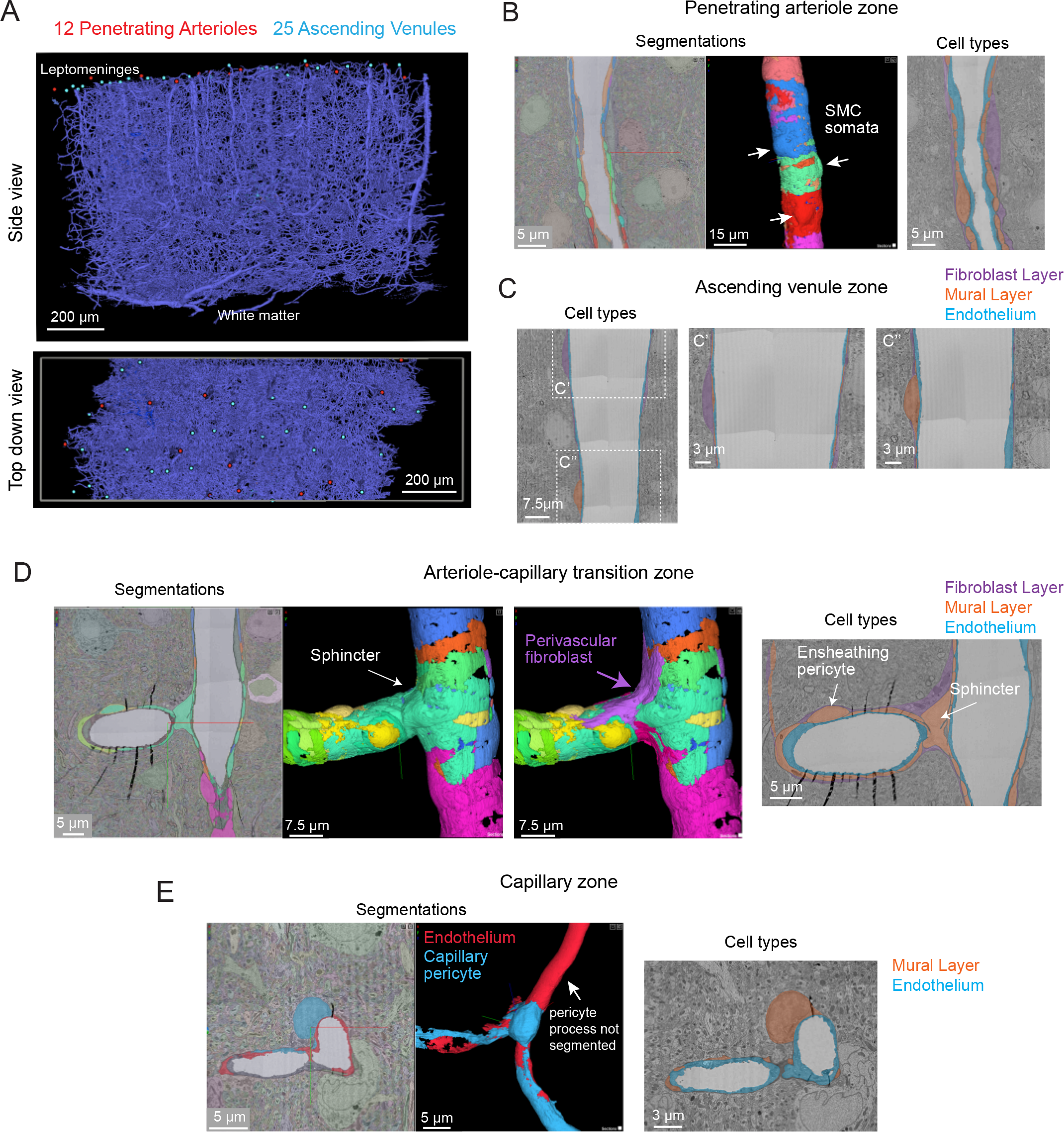
All vascular zones are represented in the MICrONS Cortical MM^3 data set. **A.** The vasculature captured in the MICrONS Cortical MM^3 data set includes 12 penetrating arterioles, 25 ascending venules, and extensive intervening capillary networks. The bottom panel shows a top-down view of the volume and reveals the originating position of penetrating vessels at the pial surface. Link to annotations: https://ngl.microns-explorer.org/#!gs://microns-static-links/mm3/vascular_annotation.json. **B.** A penetrating arteriole with some cell types within the vascular wall segmented. Here, the rings of smooth muscle cells are selected for 3D viewing (arrows). On the right, the three distinct cell types that make up the vascular wall (endothelial cells, mural cells, and perivascular fibroblasts) are shown through manual labeling. MICrONS Cortical MM^3 x,y,z coordinates at: 225166, 107266, 18188. **C.** An ascending venule exhibits a vascular wall that is thinner than the arteriole, but still contains endothelial cells, mural cells, and perivascular fibroblasts. These cell types were not well separated by segmentation and therefore only manual labeling of these cell layers is shown. Coordinates at: 262168, 104664, 24791. **D.** The arteriole-capillary transition zone is a small segment of vasculature that intervenes between the penetrating arteriole and true capillary bed. The endothelium, mural cells (sphincter cell and ensheathing pericyte), and perivascular fibroblasts are generally segmented in this region of interest. Here, we show the mural cell layer (left and middle) which involved cobbling of many separately segmented structures to visualize the overall morphology of the sphincter cell and ensheathing pericyte. The sphincter cell was classified based on prior descriptions by Grubb *et al*. showing a cuff-like cell at the penetrating arteriole just before branching into the transitional zone, with local decrease in vessel diameter. On the right, we add a perivascular fibroblast, which may not have completely captured in entirety but shows feasibility of its separate segmentation. On the right, the location of different cell layers is shown through manual labeling. Coordinates at: 168985, 165246, 19171. **E.** The capillary wall consists of only endothelial cells and pericytes. These two cells are sometimes separately segmented, as in the case shown here. However, it is not consistent throughout the network. One process of the pericyte is segmented together with the endothelium. On the right, the location of these cells is shown through manual labeling. Note the fibroblast layer is absent in capillaries. Coordinates at: 231661, 104310, 24813.

Unlike the MICrONS Layer 2/3 data set, endothelial and mural cells have been segmented separately in Cortical MM^3. However, these segmentations are not consistent throughout the microvasculature. The segmented endothelium is not of individual cells, but of most of the endothelial layer. Further, mural cells are complex in structure and an individually segmented region may represent fragments of cells. Nevertheless, they demonstrate feasibility of segmenting vascular cells and provide biological insight. For example, smooth muscle cells of the penetrating arteriole viewed in 3D reveal the position of their somata, which bulge slightly from the vascular wall (**Fig. 7B, left, middle**). The tiling of their individual territories and details of their circumferential processes can be examined. In an accompanying 2D image we verified the positions of different cell types along the same arteriole (**Fig. 7B, right)**. In contrast to the arteriole, the wall of ascending venules is much thinner, and as such, the segmentation of endothelial and mural cells along venules examined is less accurate than arterioles. We provide only labeling in 2D to show how vascular cells are positioned along the venular wall (**Fig. 7C**).

The extensive microvascular network could be seen throughout the data set, including capillaries and the transitional zones bridging the capillaries to arterioles and venules. In one example, a small penetrating arteriole extends an offshoot toward the capillary network (**Fig. 7D**). A pre-capillary sphincter cell (a presumed mural cell type) is seen as a cuff around the origin of this offshoot where the vessel diameter is locally decreased (Grubb et al., 2020). Sphincter cells are morphologically distinct from pericytes and exhibit high reactivity to vasoactive mediators during neurovascular coupling (Zambach et al., 2021), as well as aberrant contraction during pathology (Khennouf et al., 2018).

The protruding soma of an ensheathing pericyte is visible just downstream of the sphincter (**Fig. 7D**). Unlike the more prevalent capillary pericytes found deeper in the capillary network, and as described for the Layer 2/3 data, this pericyte subtype of the transitional zone exhibits longer processes than SMCs but shorter than those of capillary pericytes (Hartmann et al., 2021a;Hartmann et al., 2021b). However, the processes completely cover the vascular circumference with fine interdigitated projections (Grubb et al., 2020). Critically, in the image shown (**Fig. 7D, middle)**, we suspect that the complexity of this cell is not captured by a single segmented volume. Rather, multiple segmented regions were cobbled together to obtain a general view of the cell coverage of the transitional segment. Ensheathing pericytes are reactive to vasoactive stimuli and the regions they cover dilate rapidly during functional hyperemia (Hall et al., 2014).

As described in the MICrONS Layer 2/3 data, capillary pericytes exhibit protruding somata and long processes that incompletely cover the circumference of endothelium (Grant et al., 2017). Pericytes of this morphology could be discerned throughout the extensive capillary beds of the sample. Pericyte and endothelial cells are occasionally segmented as individual cell types along the capillary wall (**Fig. 7E**). However, the segmentation of individual pericytes may be incomplete, as evidenced by a missing process in the 3D view shown. Throughout the capillary network, the nuclei of pericytes and endothelial exhibited clear contrast, providing a unique opportunity to understand pericyte-endothelial positioning and arrangement in the microvasculature. This information may add an additional parameter to consider for in silico studies of capillary flow through cortical vascular networks.

#### Perivascular fibroblasts

Perivascular fibroblasts lie just abluminal to the mural cells on larger parenchymal vessels in the brain (Bonney et al., 2021). However, they are absent in the capillary zone. The physiological role of perivascular fibroblasts remains largely unknown. Single cell transcriptomic analyses of brain vasculature have shed light on their gene expression profiles, which suggest potential roles in basement membrane protein production and contribution to cerebrovascular structure (Vanlandewijck et al., 2018). In some pathological scenarios, such as neurodegenerative disease, ischemia and injury, perivascular fibroblasts proliferate to form scar tissue (Soderblom et al., 2013). Consistent with prior work, we observed perivascular fibroblasts in arteriole, arteriole-capillary transition (**Fig. 7B,D**), but not in capillaries (**Fig. 7E**). They are also present on venules (**Fig. 7C**), but only those of larger diameter and more superficial of the brain. Morphologically, perivascular fibroblasts exhibit flattened somata and thin lamellar processes that cover the vessel surface.

#### Perivascular macrophages

Perivascular macrophages are resident immune cells of the brain with close association to the vasculature. They have been shown to communicate with other neurovascular cell types, are a source of reactive oxygen species, and contribute to a variety of disease processes (Faraco et al., 2016). They occupy the same perivascular space as fibroblasts and may also exhibit morphological similarities. Thus, detailed examination will be required to understand how to separate these two cell types at the ultrastructural level. Perivascular macrophages are highly phagocytic and should presumably be distinguishable from fibroblasts by a higher density of electron dense lysosomes.

#### Subcellular compartments

Image contrast in the Cortical MM^3 data was sufficient to resolve subcellular features (endothelial cell junctions, caveolae, Golgi cisternae) (**Fig. 8**). Instances of caveolar fusion with the endothelial plasma membrane were observed (**Fig. 8B, Bi**).

**Figure 8.**
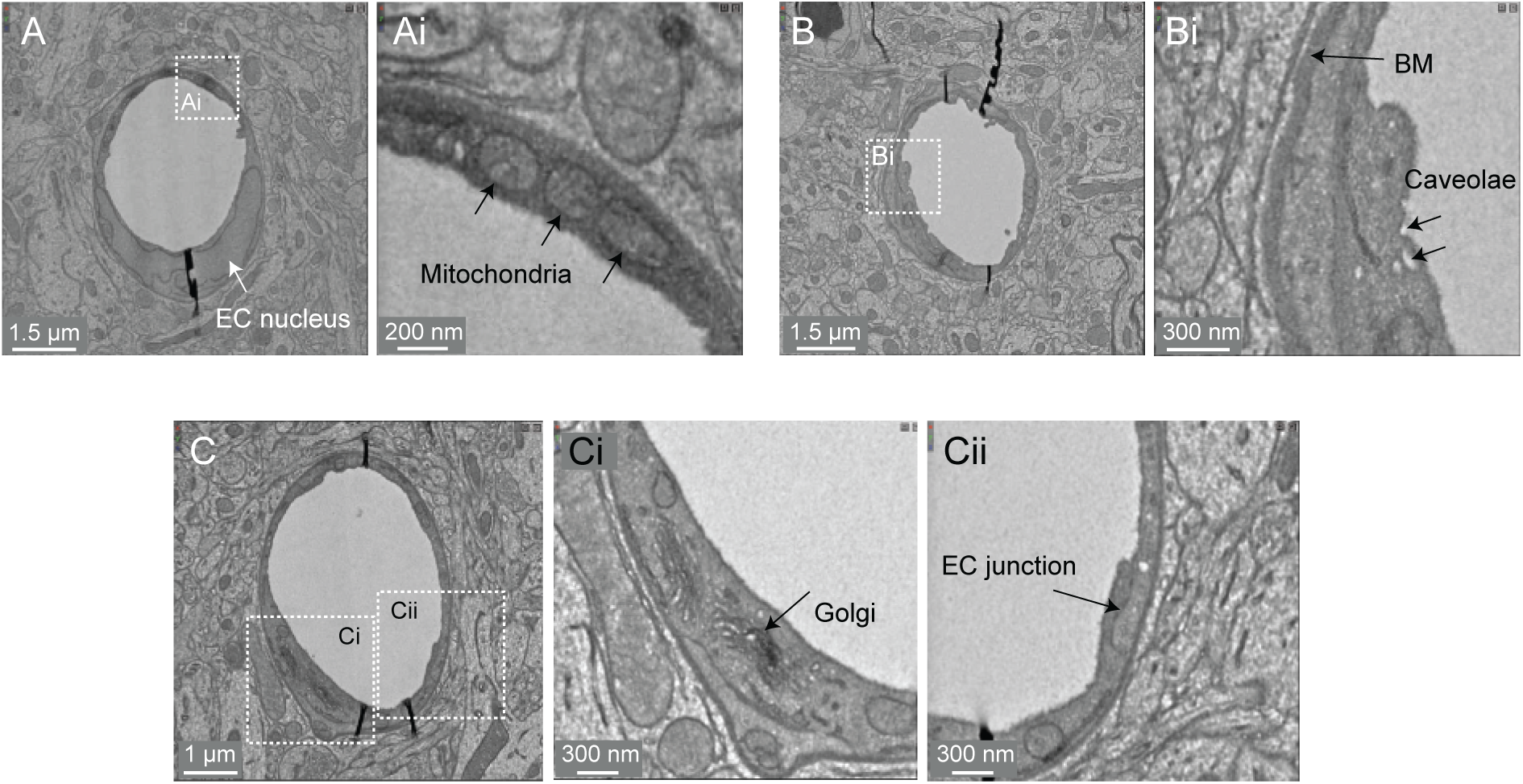
Subcellular features of the endothelium in the MICrONS Cortical MM^3 data. **A.** Capillary cross-section with inset (Ai) showing mitochondria. MICrONS Cortical MM^3 x,y,z coordinates at: 141179, 177334, 21310. **B.** Capillary cross-section with inset showing endothelial caveolae and basement membrane. Coordinates at: 167382, 134731, 20338. **C.** Capillary cross-section with inset (Ci) showing Golgi cisternae and separate inset (Cii) showing endothelial junction. Coordinates at: 129596, 117071, 20179.

#### Perivascular space

The perivascular space is a fluid filled space surrounding larger brain vessels that is vital for influx and efflux of cerebrospinal fluid in the clearance of metabolic waste from brain tissue (Iliff et al., 2012;Wardlaw et al., 2020). However, there remains a lack of clarity on the anatomy of the perivascular spaces, leading to controversy in pathways for CSF flow and where they reside in the vascular wall (Bakker et al., 2016). 3D-EM data may provide an opportunity to define the architecture of the perivascular space. Details including the vessel types that exhibit a perivascular space (arterioles vs. venules, small vs. large diameter vessels), the diameter/volume of these spaces, and the neurovascular cell types forming their boundaries could be examined in detail. However, caution is needed to ensure that these spaces are adequately preserved in fixed tissue specimens.

In the Cortical MM^3 data, perivascular spaces could be observed primarily around larger penetrating arterioles and ascending venules, and they were most apparent in the upper layers of cortex (**Fig. 9A**). The perivascular space diminishes as the vessels become smaller in diameter (**Fig. 9Ai**). The space eventually becomes continuous with the perivascular fibroblast layer, flanked by two layers of basement membrane (**Fig. 9Aii, Aiii**). Smaller arterioles, arteriole-capillary transition zones, and capillaries did not exhibit overt perivascular spaces though they were encased in the basement membrane. This raises the question of whether CSF flows along the walls of the small vessels and capillaries, and if so whether it occurs within the vascular basement membrane. It also suggests that perivascular fibroblasts may be involved in adherence of the vascular basement membranes, and in doing so influence the extent of the perivascular space.

**Figure 9.**
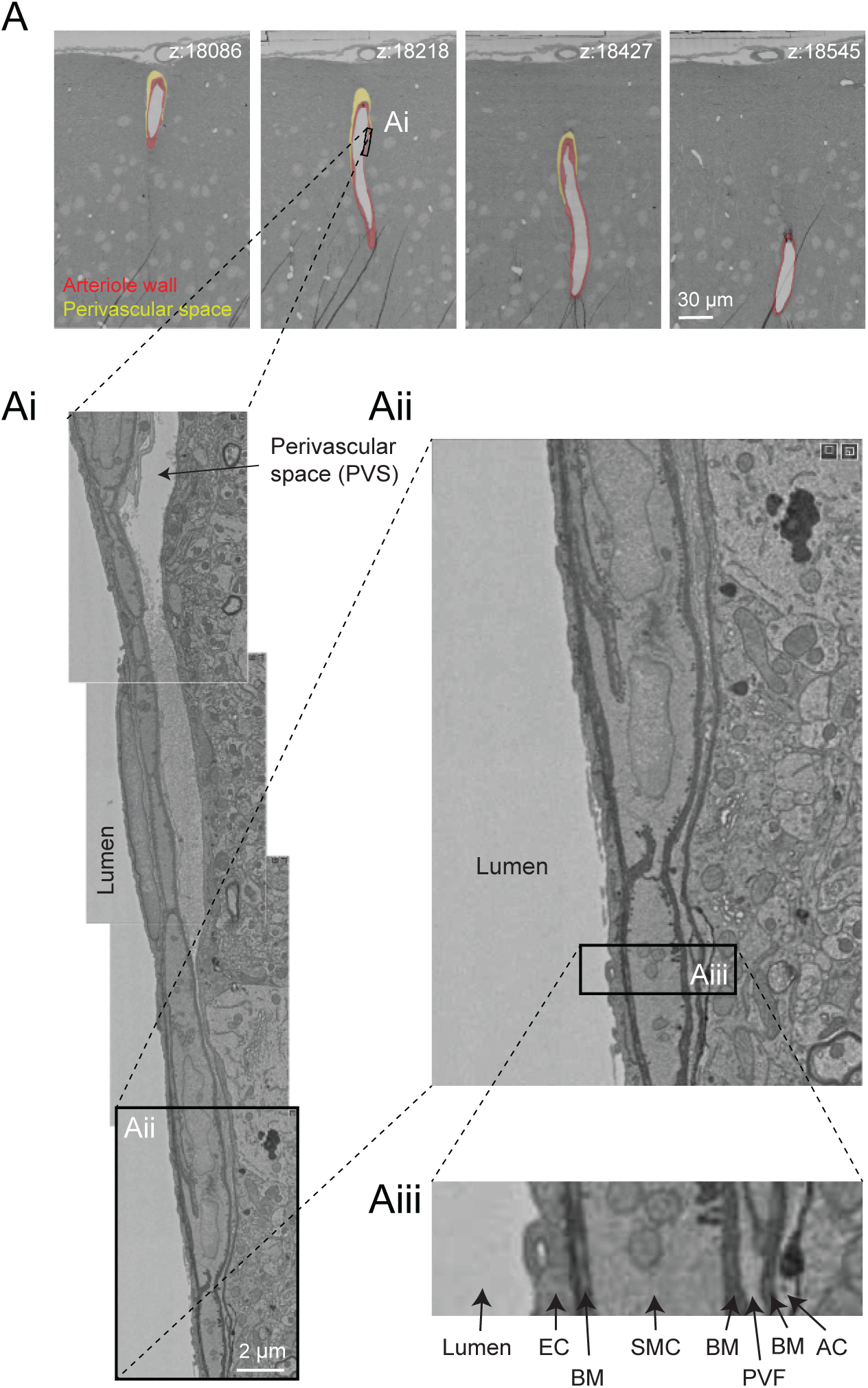
Perivascular space and layers of the arteriole wall. **A.** A large penetrating arteriole in the MICrONS Cortical MM^3 data set exhibits a clear perivascular space. **Ai.** Magnified view of the arteriole wall showing the perivascular space diminishing in size with greater cortical depth. **Aii, Aiii.** Magnified view showing layers of the arteriole wall, revealing that the perivascular space becomes continuous with the perivascular fibroblast layer as the perivascular space diminishes. EC - endothelial cell, BM - basement membrane, SMC - smooth muscle cell, PVF - perivascular fibroblast, AC - astrocyte. Coordinates of the penetrating arteriole: 175231, 117475, 19297.

#### White matter

The Cortical MM^3 data set contains white matter just below the cortex, where microvasculature is seen alongside densely packed myelinated axons (**Fig. 10A,B**). There is sufficient contrast to discern the boundaries of endothelial cells and mural cells in white matter, as well as subcellular features, such as caveolae. The interaction of white matter astrocytes with the vascular wall may be accessible to study, as some of the white matter astrocytes have been segmented (**Fig. 10C**). The vasculature in white matter also contains one branch of a principal cortical venule, which is a large ascending venule that serves as the primary drainage route for juxtacortical white matter (**Fig. 10D**)(Duvernoy et al., 1981).

**Figure 10.**
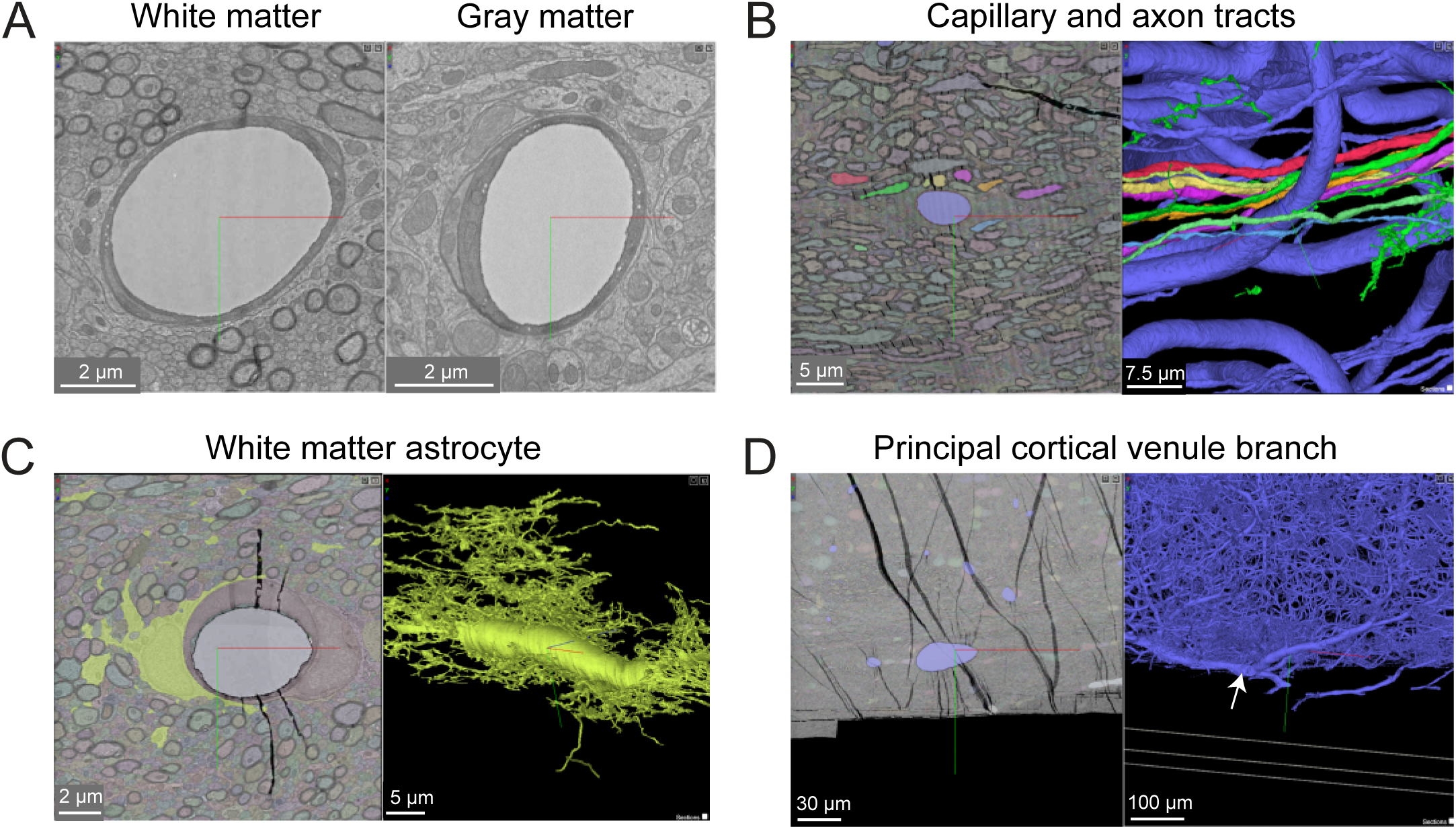
White matter vessels in MICrONS Cortical MM^3 data. **A.** Representative cross section of capillary-sized vessels in white matter (left) and gray matter (right). White matter capillary cross-section x,y,z coordinates at: 114029, 277354, 17647. Gray matter vessel cross-section x,y,z coordinates at: 109231, 208260, 17714. **B.** A white matter capillary embedded within the axon tracts of the white matter (left). In 3D view, the vascular lumen segmentation is displayed, alongside select axons that have been segmented. Coordinates at: 155004, 269113, 21080. **C.** A perivascular white matter astrocyte. In 3D view, the coverage of the capillary wall can be seen. Coordinates at: 150176, 279591, 18383. **D.** A branch of a deep draining venule, principal cortical venule, is present within the data set. Coordinates at: 195156, 273647, 18192.

This gray-white matter sampling can help to clarify differences in composition and subcellular features of the neurovascular unit as blood vessels course into the white matter environment. For example, it should be possible to understand structural differences in pericyte coverage and association with the endothelium, as well as the ratio of pericytes and endothelial cells per capillary length. Given the vulnerability of white matter to ischemia and in neurodegenerative diseases, understanding how the neurovascular unit differs between gray and white matter will be of great importance.

#### Accompanying physiological data

Remarkably, the 3D-EM data produced by the MICrONS consortium has accompanying multiphoton imaging data on neuronal activity from the same tissues, where the microvasculature served as a fiduciary to co-register *in vivo* and *post-mortem* data. This physiological data includes neuronal calcium dynamics during visual stimulation, and separately, vascular structure (intravenous fluorescent dye). This is proof-of-principle that ultrastructural features of the microvasculature may eventually be linked to neurovascular dynamics captured *antemortem*.

### Additional studies of mouse visual cortex

A 2016 study by Lee *et al*. provided a sample from visual cortex layer 1 and layer 2/3 with a volume of 450 x 450 x 150 μm (Lee et al., 2016). Image contrast was high, allowing cellular and non-cellular (basement membrane) components of the vascular wall to be readily discerned. The volume contains a portion of a penetrating arteriole and surrounding capillary networks. However, the vascular-specific compartments (vascular lumen, wall, astrocytes) are not segmented.

### Hippocampus

The hippocampus has a different vascular supply and microvascular architecture compared to the cerebral cortex, and these differences may underlie vulnerabilities to blood flow insufficiency (Shaw et al., 2021). Thus, comparative 3D-EM studies across different brain regions will be valuable in future studies. A study by Bloss *et al*. imaged a volume of mouse hippocampal CA1 region spanning stratum radiatum and stratum lacunosum-moleculare (Bloss et al., 2018). The sample is 350 x 200 x 17 µm in size and contains portions of a venule network and some surrounding capillaries. The image contrast is excellent, allowing a clear separation of vascular cell types, subcellular structures, and the basement membrane layers (**Fig. 11A**). Curiously, some vessels exhibit enlarged astrocytic endfeet not seen in the cortical data sets described above (**Fig. 11B**). Thin projections of the vascular basement membrane also extended into the parenchyma. Whether these features are true distinctions of hippocampal versus cortical vasculature or consequences of tissue preparation remain to be clarified.

**Figure 11.**
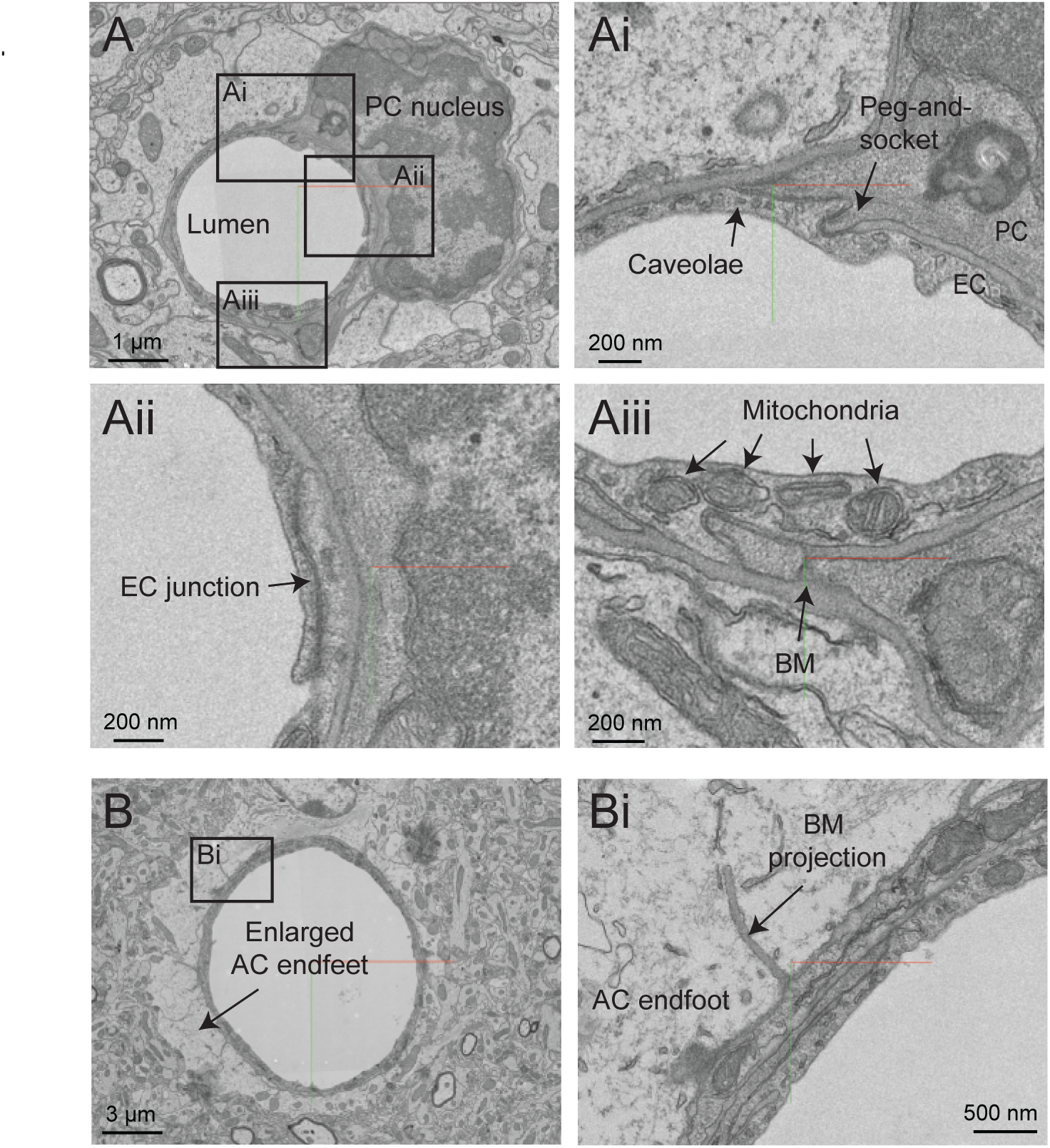
Ultrastructural view of microvasculature in the mouse hippocampus. **A.** Representative images from data of Bloss *et al*. taken from mouse hippocampus. The image shows a capillary cross-section where a pericyte soma is located. Insets from Ai to Aiii show a variety of subcellular structures that can be visualized. Coordinates at: 20865, 54458, 146. **B.** A separate capillary cross-section taken from the same volume exhibits enlarged astrocytic endfeet against the vascular wall. An inset (Bi) shows a thin projection of the basement membrane extending abluminally. Coordinates at: 25774, 59225, 86.

### Finch basal ganglia

Zebra finch is a widely used model system in behavioral research, particularly with respect to song development and auditory processing. Two zebra finch brain data sets (j0126 and j0251, basal ganglia, Area X) were imaged with high contrast 3D-EM and show many details of the cellular and subcellular structure of the avian vasculature (**Fig. 12A**). The smaller j0126 data set (97.9 × 95.6 × 115 μm) contains a venule and surrounding capillaries. Close inspection of the capillaries revealed similar arrangement of endothelial cells and pericytes as seen in the mouse brain. Pericyte and endothelial interdigitations were widely observed, and in some cases pericyte pegs contacted the endothelial nucleus as reported previously in mouse (Ornelas et al., 2021). Interestingly, in the larger j0251 data set (256 × 256 × 384 μm), pericyte projections were seen to bypass the endothelial layer and protrude into the luminal space and end in a small cilia-like structure (**Fig. 12B-Biii**), opening the possibility that pericytes use these projections for direct sensation of blood flow. This is a feature that we have thus far not seen in the mouse brain.

**Figure 12.**
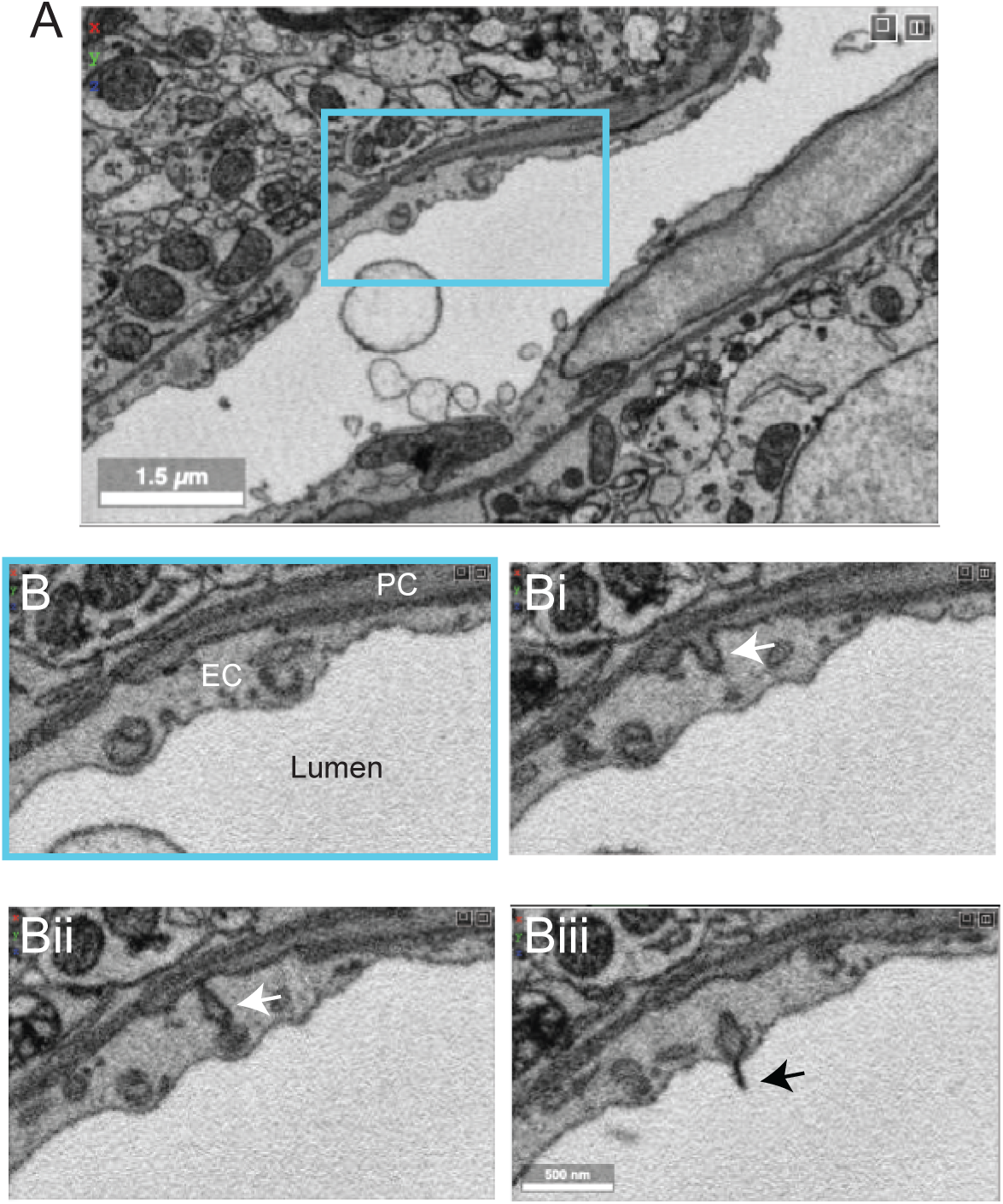
Pericyte projection into vessel lumen in finch brain capillaries. **A.** A capillary from finch basal ganglia (data set: j0251), located at x,y,z coordinates: 19129, 12810, 7330. **B-Biii.** Magnified view of inset in panel A showing a pericyte projecting into the luminal space and ending in a cilia-like structure. The four panels span 10 slices in the z dimension, corresponding to ∼200 nm of distance.

### Human cortex

A recent study from Shapson-Coe *et al*. examined a small fragment of human cortex (temporal lobe), resected from a patient with drug-resistant epilepsy during surgery to remove an epileptic focus (Shapson-Coe et al., 2021). Immediately after excision, the sample was immersion-fixed overnight. The hippocampus was sclerotic and exhibited signs of pathology consistent with epilepsy, but the resected cortical sample appeared normal on traditional histopathology.

The sample spans ∼3 mm at its longest width, ∼2 mm at its longest height, and ∼0.15 mm in thickness. It includes sections of parenchymal arterioles and venules and surrounding capillary networks (**Fig. 13A**). Despite the vascular lumen being mostly collapsed, the nuclei of vascular cells could be categorized as mural cells or pericytes based on distance from the lumen and position relative to the endothelial and parenchymal basement membranes (**Fig. 13B**). A total vascular length of 22.6 cm was segmented, in which the authors detected ∼8k vascular cells (4604 endothelial cells, 3549 pericytes, 574 SMCs and 79 fibroblast-like cells). The astrocyte endfeet surrounding the vasculature were swollen, which is unlike that seen in the samples prepared from mouse or finch and may be due to the limitations of *post-mortem* sample collection (**Fig. 13B**).

**Figure 13.**
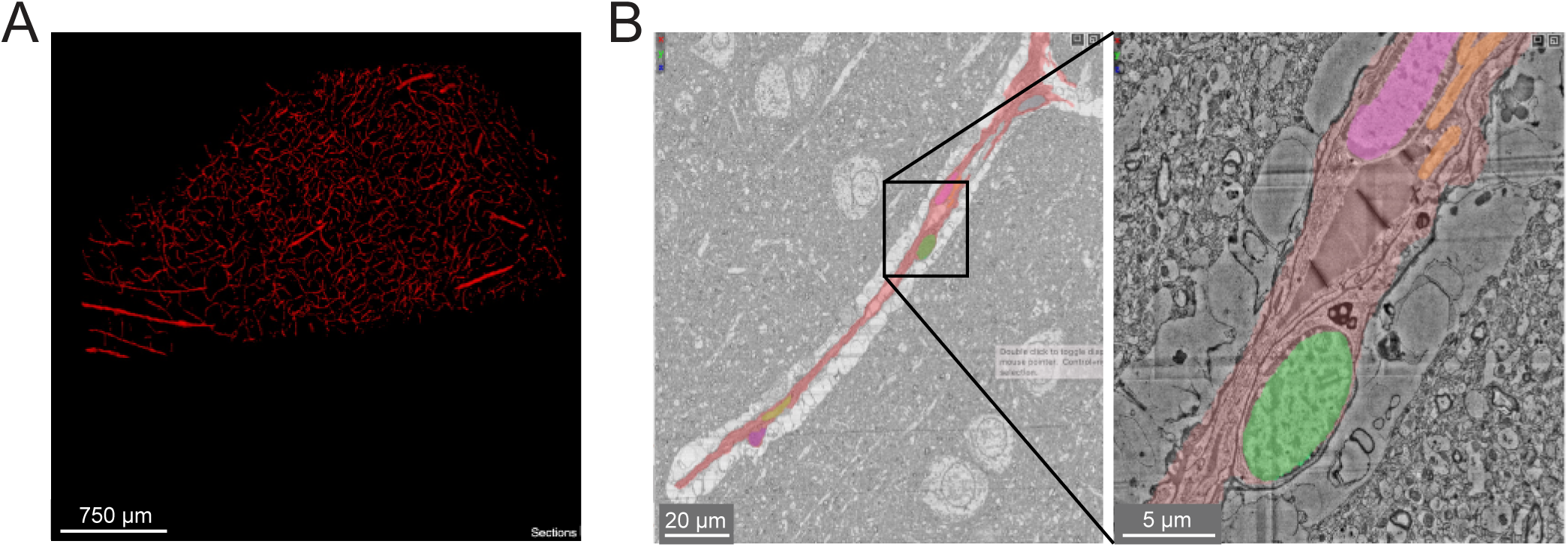
Human cerebral cortex microvasculature. **A.** Broad view of the microvasculature captured in the data set of Shapson-Coe *et al*. **B.** A capillary-sized vessel showing general segmentation of the vasculature. The nuclei of vascular cells were separately segmented into pericytes or endothelial cells, based on their location relative to the blood vessel lumen and basement membrane layers, as well as their appearance in EM. In the example shown, the pericyte nucleus is green and the endothelial nucleus is pink. Left image x,y,z coordinates at: 297270, 74996, 5229. Right image x,y,z coordinates at: 299348, 73663, 5229.

A notable feature unique to this human tissue sample is the presence of many perivascular oligodendrocytes that lined the vascular wall, particularly the radially-oriented vessels of white matter. This may be why oligodendrocytes are a prevalent cell type isolated from vascular-enriched single cell transcriptomic studies (Yang et al., 2022). Another observation was the existence of string capillaries, which are basement membrane tubes with pericytes encased, but no endothelial cells. Their occurrence is not uncommon in the normal brain (Brown, 2010), but prevalence can increase in pathological context leading to reduced capillary connectivity and blood flow impairment (Challa et al., 2004).

## DISCUSSION

We surveyed the vasculature in 6 recent large-scale 3D-EM studies (**Table 1**). These data spanned different species (mouse, finch, human) and brain regions (cortex, callosum, hippocampus, basal ganglia). The data sets captured images across different zones of the microvasculature (arterioles, capillaries, venules and transitional segments), and were of sufficient quality to visualize subcellular features relevant to blood-brain barrier function (endothelial junctions, caveolae, basement membrane, astrocytic coverage), blood flow (positioning of endothelial and pericyte nuclei, mural cell coverage, endothelial microvilli, microglial-vascular interaction), bioenergetics (mitochondria), cellular communication (caveolae, peg-and-socket interactions) and cerebrospinal fluid flow (perivascular space). The biological insight that can be garnered from these data is immense. Further work is needed to segment, annotate and quantify cerebrovascular features. These efforts will stimulate the development of new hypotheses, design of novel physiological studies, and improved *in silico* modeling approaches.

To fully leverage 3D-EM for vascular research, it will be necessary to accurately segment the cellular components of the vascular wall: endothelial cells, mural cells (pericytes and smooth muscle cells), perivascular fibroblasts, and perivascular macrophages. Together with already segmented parenchymal components (neurons, astrocytes, microglia, and oligodendrocytes), this would account for all cells that comprise the neurovascular unit, a framework used to understand how neurons and other cell types communicate with nearby vessels to serve metabolic needs of the brain (Schaeffer and Iadecola, 2021). Advances in the MICrONS Cortical MM^3 data set have shown feasibility of segmenting individual cell types of the vascular wall. However, further work is needed to improve accuracy and consistency of the current segmentations and to identify the specific challenges posed by vascular cells. In addition, existing data sets offer the possibility to identify and map innervating neurons of cerebral vessels, providing insights into flow regulation by direct release of neurotransmitter on the vascular wall (Hamel, 2006).

Mapping the prevalence and distribution of vascular ultrastructures is essential for our understanding of vascular functions across different zones. For example, the nuclei of endothelial cells and pericytes are easily distinguished in the MICrONS Cortical MM^3 data. How the nuclei localize in the network and impinge upon the luminal space, may hold insight into how blood flow is distributed through complex capillary networks (Østergaard et al., 2014;Schmid et al., 2017). Further, it is likely that endothelial microvilli contribute to blood flow resistance, and conceivably sense blood flow to convey signals to the endothelium and mural cells. Therefore, understanding their distribution at the level of individual capillary segments would be valuable.

Mapping endothelial junctions and caveolae on a large scale would generate an unprecedented 3D map of blood-brain barrier structure. Endothelial junctions and caveolae bear similarity to the post-synaptic density and pre-synaptic vesicles of neuronal synapses, respectively. Thus, existing machine learning algorithms are likely adaptable for detection of these vascular features. Identification of endothelial junctions will also reveal the boundaries of individual endothelial cells as they tile together to form the inner layer of the vascular tube. Additionally, mitochondria in vascular cells have a similar appearance to those already segmented in parenchymal cells (see MICrONS Layer 2/3). Mapping the density of mitochondria across the vasculature may reveal where energy-demanding processes, such as blood-brain barrier transport or maintenance of membrane potential, are most active.

The ability to manually annotate, and therefore count and quantify, the number of cellular/subcellular structures already provides significant insight on vascular structure. For example, we have annotated endothelial and pericyte nuclei, and traced pericyte processes and endothelial junctions. In prior work, basic distance measurements were performed in Neuroglancer allowing extraction of quantitative data on peg-and-socket contacts between endothelial cells and pericytes (Ornelas et al., 2021) These endeavors have only scratched the surface of information within these volumes, and there are no barriers to further exploration in this fashion, except time. Further, Neuroglancer is one of many browsers for exploration of volume imaging data, and others including webknossos, knossos, pyknossos, napari, bigdataviewer, may be more versatile for annotation. Thus, the availability of 3D-EM data through a variety of online tools will maximize their use by the scientific community.

A potential limitation and topic for further investigation is whether the parameters of tissue collection used in the current data sets (anesthetic, *post-mortem* interval, fixative, perfusion pressure, etc.) adequately preserved the native structure of the vascular lumen, vascular cell types, and perivascular space. The shape of the vessel wall can be easily distorted by contraction of mural cells or loss of intraluminal pressure, making quality of tissue preparation of utmost importance. It is already known that the choice of fixation approach greatly influences the size of the extracellular space (Cragg, 1980;Pallotto et al., 2015) and this may result in disparate outcomes for both neuronal and vascular structures alike. For example, brains prepared with aldehyde-based fixative for EM show nearly full coverage of blood vessels with astrocyte endfeet (Mathiisen et al., 2010) while only two-third coverage is seen with cryo-EM (Korogod et al., 2015). One way to optimize tissue preparation is to image the same regions both *in vivo* and *post-mortem*, as done in the MICrONS pipeline. The *in vivo* two-photon imaging data may then be compared to the 3D-EM data to understand how well vascular diameter and wall structure is preserved. Further, cautionary statements should be added to publications when there are signs of poor vascular preservation, include stacking of red blood cells in arterioles, wide-spread swelling of astrocytes, and ruffling of endothelial cells.

The work and costs involved in creating and analyzing large-scale 3D-EM on the scale of the MICrONS MM^3 data are extensive. It is not readily accessible to smaller, individual laboratories seeking to address specific questions in their studies. However, some labs may wish to generate more manageable 3D data sets for manual segmentation and annotation. The challenge is that vasculature is a relatively sparse in brain tissue compared to neurons, and one must know where to collect smaller data sets. A way to overcome this challenge is to first survey the broader vascular network with wider field imaging, and then perform high-resolution imaging specific regions of interest (Guérin et al., 2019). The public resources described here can serve as valuable guides to decide where to collect vascular data for more targeted studies. Further, techniques such as X ray micro CT angiography (Ghanavati et al., 2014) could be used to survey vasculature across the whole brain prior to EM-level imaging within distinct brain regions.

## Author contributions

AYS and AK conceptualized the study. SKB, VCS, SFH, and AK surveyed the data to find notable vascular features and contributed to preparation of figures. JK collected and provided access to finch data sets. MT provided insight into navigation and presentation of MICrONS consortium data. AYS wrote the manuscript with contributions and feedback from all authors.

## Acknowledgements

The authors thanks F. Collman, and N. da Costa for helpful discussions.

